# Piezo-dependent surveillance of matrix stiffness generates transient cells that repair the basement membrane

**DOI:** 10.1101/2023.12.22.573147

**Authors:** Aubrie M. Stricker, M. Shane Hutson, Andrea Page-McCaw

## Abstract

Basement membranes are extracellular matrix sheets separating tissue layers and providing mechanical support, and Collagen IV (Col4) is their most abundant protein. Although basement membranes are repaired after damage, little is known about repair, including whether and how damage is detected, what cells repair the damage, and how repair is controlled to avoid fibrosis. Using the intestinal basement membrane of adult *Drosophila* as a model, we show that after basement membrane damage, there is a sharp increase in enteroblasts transiently expressing Col4, or “matrix mender” cells. Enteroblast-derived Col4 is specifically required for matrix repair. The increase in matrix mender cells requires the mechanosensitive ion channel Piezo, expressed in intestinal stem cells. Matrix menders are induced by the loss of matrix stiffness, as specifically inhibiting Col4 crosslinking is sufficient for Piezo-dependent induction of matrix mender cells. Our data suggest that epithelial stem cells control basement membrane integrity by monitoring stiffness.

## Introduction

Extracellular matrix (ECM) is an essential conserved component of all animal tissues. In addition to providing structural support, ECM influences cell adhesion, shape, signaling, and behaviors. When tissues are damaged, ECM needs to be repaired, and pathological ECM damage is a feature of many human diseases^1,2^. Despite the critical importance of ECM repair, little is known about the cellular processes and molecular signals that orchestrate repair. At the most fundamental level, it is unknown whether and how cells detect ECM damage. Moreover, ECM proteins, including collagens, are among the most stable proteins in the body, with the longest-lived exhibiting half-lives on the order of an animal’s lifetime^3–6^. ECM longevity presents a challenge, as repairing with excess ECM causes fibrosis. In this study, we identify a mechanism that cells use to sense damaged matrix, and they respond by transiently producing matrix until repair is completed.

ECM can be subdivided into two types, stromal matrix and basement membrane, the focus of this study. Basement membranes are thin sheets that underlie epithelia and wrap around muscles and organs. Their most abundant protein is Collagen IV (Col4), a heterotrimer that self-assembles *in vitro* into networks and provides structural support^7^. Self-assembly raises questions about the nature of basement membrane repair. Is there a continuing self-assembly process for maintenance that also functions passively to repair damage? Or is repair an active process, with cells sensing damage and directing repair? Basement membranes are ancient, arising along with multicellularity itself^8^. Given the evolutionary conservation, *Drosophila* offers an excellent model to address questions of basement membrane homeostasis and repair. Flies have the same basement membrane structure and composition as mammals, with all the main components but fewer copies. There is only one type IV collagen (Col4) heterotrimer composed of the two collagen subunits Col4a1 and Col4a2 (aka Viking, Vkg); there are two laminin heterotrimers, one nidogen gene, and one perlecan gene^9^. Functional GFP- tagged alleles of Vkg offer tremendously powerful reagents for imaging and genetics^10^. We use the *Drosophila* posterior midgut as a model to analyze basement membrane repair. The fly gut is a monolayer epithelium derived from stem cells that generate two lineages, enterocytes and enteroendocrine cells, with intermediates in between^11–13^.

Between the epithelium and the surrounding peristalsis muscles there is basement membrane, and it is continuous with the basement membrane enveloping the muscles (Fig. 1A). We damage the basement membrane by feeding the flies dextran sodium sulfate (DSS), which gets visibly lodged in the gut basement membrane, fractures its structure, and reduces its stiffness^14^. After stopping DSS feeding, basement membrane repair occurs within two days, providing a temporal window for analyzing repair^14^.

**Figure 1.**
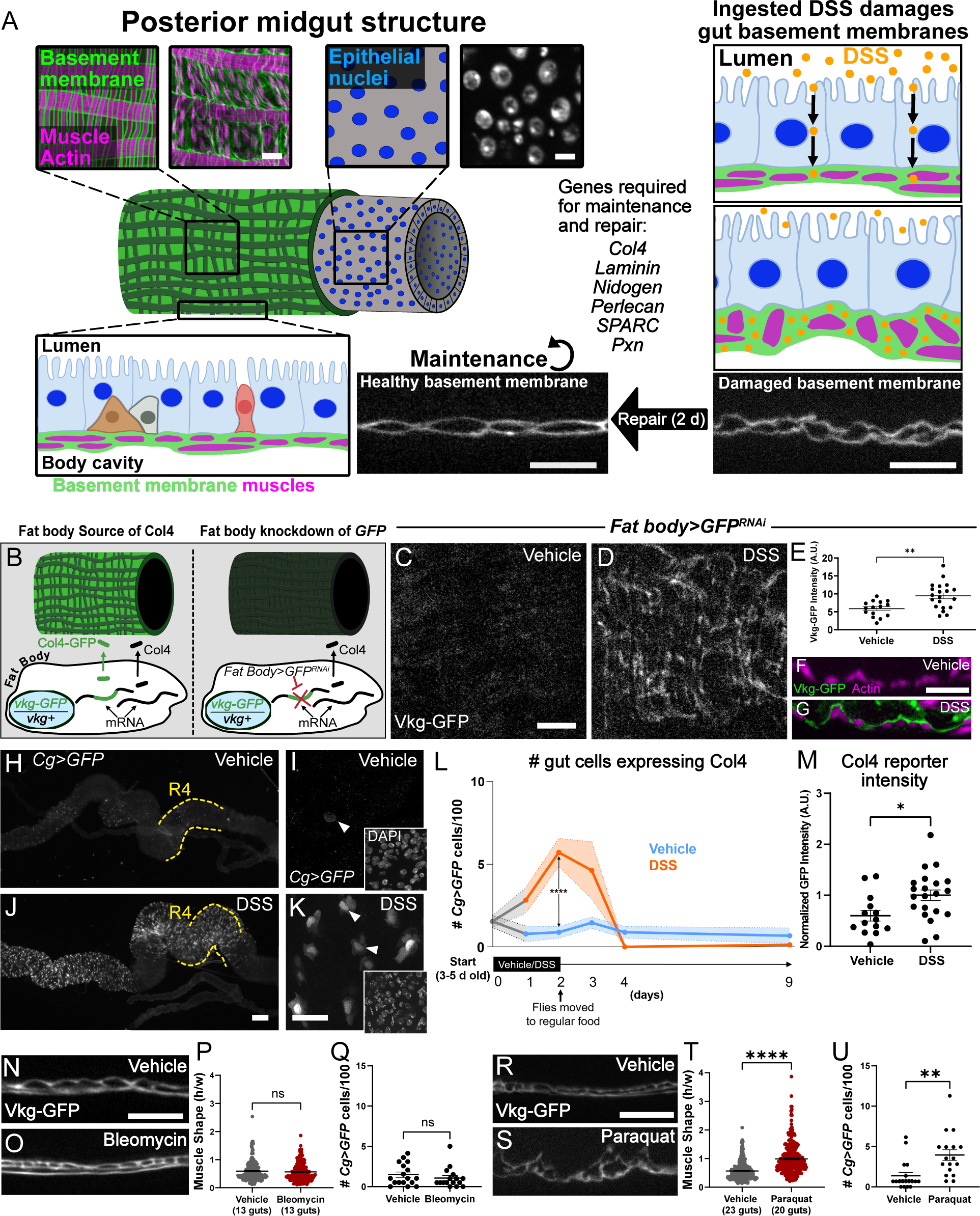
“Matrix mender” gut cells upregulate Col4 after basement membrane damage. **(A) Left: *Drosophila* midgut.** The gut tube is covered with peristalsis muscles (magenta) wrapped in basement membrane (green). Underneath the muscle layer, the gut monolayer epithelium (blue nuclei/DAPI) comprises absorptive enterocytes (blue), stem cells (brown), enteroblasts (gray), and enteroendocrine cells (red), all sitting on basement membrane continuous with that of the muscles (green). Scale bars, 10 µm. **Right: DSS damage.** DSS (orange dots) is ingested, transported through the enterocytes, and deposited in the underlying basement membrane where it fractures the basement membrane making it less stiff, resulting in visibly dysmorphic muscles (described in Howard et al, 2019). Basement membranes and muscle shapes are fully repaired 2 days after DSS is removed. Both repair and maintenance of the basement membrane require basement membrane components and regulators (see Fig. S1 and Howard et al, 2019, for data). Scale bars, 10 µm. **(B-G)** Collagen IV (Vkg) in the gut basement membrane comes from the fat body in undamaged conditions. B depicts the experiment knocking down GFP of Vkg-GFP specifically in the fat body with *c564-Gal4*, data in C (*en face* view). After DSS damage, some gut Vkg-GFP came from another source (D,E). Each dot in E represents a gut. Cross-sections (F,G) show the new Vkg source deposits Vkg-GFP between the epithelium and the muscles. Full data set for fat-body knockdown and the specificity of the *c564-Gal4* driver shown in Fig. S2. **(H-K)** Col4 reporter *Cg>GFP* identified unknown cells that respond to DSS in the gut. Anterior (left), posterior (right). Posterior R4 midgut is the focus of this study. Scale bar, 100 µm in H,J, scale bar, 20 µm in I,K. **(L-M)** Collagen-expressing cells (“matrix menders”) increased in number during basement membrane damage, then decreased during repair (L). Shaded regions represent SEM. 5-21 guts analyzed per timepoint and treatment, see Fig. S3A for dataset. Significance by *t*-test. In addition to number, expression level increased after DSS (M), see arrowheads in I,K. Each dot represents a gut. Mean +/- SEM, significance by *t*-test. **(N-Q)** Feeding bleomycin did not damage the basement membrane, as indicated by muscle shape (N-P), even though it did cause DNA damage and stem cell proliferation (see Fig. S3B,C). Bleomycin did not induce *Cg>GFP+* matrix menders (Q). Mean +/- SEM, significance by *t*-test. Scale bar, 10 µm. **(R-U)** Feeding paraquat damaged the basement membrane, indicated by changes in muscle shape aspect ratio (R-T). Paraquat induced *Cg>GFP*+ matrix menders (U). Mean +/- SEM, significance by *t*-test. Scale bar, 10 µm.

In this study, we find that genes encoding all major basement membrane components are required for both repair and maintenance. However, after damage, the source of Col4 for the midgut changes: without damage, Col4 originates from the fat body, but after damage, Col4 is transiently expressed locally by gut epithelial cells we call “matrix menders,” which are specialized short-lived enteroblasts. Knocking down Col4 in enteroblasts specifically inhibits gut basement membrane repair. For damage to induce matrix menders requires the mechanosensitive channel *Piezo*, expressed in neighboring intestinal stem cells (ISCs). Indeed, specifically reducing basement membrane stiffness by reducing Col4 crosslinking is sufficient to induce matrix-mender cells in a Piezo-dependent manner. Our results suggest that epithelial stem cells surveil basement membrane integrity by monitoring its stiffness, responding to defects by generating short-lived repair cells.

## Results

### Basement membrane repair and maintenance require many of the same components

DSS directly damages the posterior midgut basement membrane of *Drosophila*^14^. After ingestion, DSS is transported through enterocytes to accumulate in the adjacent basement membrane (Fig. 1A). This mechanism is different from mice, where DSS is excluded by the intact epithelium^14,15^. In flies, DSS damages basement membranes, making them visibly fractured as visualized by TEM, less dense as measured with fluorescence and super-resolution microscopy, and less stiff as measured in a tensile strain assay^14^. The basement membrane mechanical damage changes the shape of the gut peristalsis muscles, which are outlined by Col4 (Fig. 1A, right side). A similar muscle shape is observed in fly guts lacking the Col4 crosslinking enzyme Peroxidasin (Pxn)^14,16,17^, and these basement membranes also have reduced tensile stiffness^14,17,18^. In both instances, gut muscles become dysmorphic when enveloped by weakened basement membrane, and thus we can infer basement membrane stiffness from gut muscle shape, measured as aspect ratio. Importantly, two days after removing DSS and feeding flies regular food, there is full restoration of basement membrane structure and the muscle shape^14^.

Previously, we determined that repair after DSS damage requires synthesis of new basement membrane proteins, including *vkg*, *LamininB1*, and *Pxn*^14^. We asked whether other conserved basement membrane components were required for repair, targeting Perlecan (*trol*), Nidogen (*Ndg*), and the Col4 extracellular chaperone, *SPARC*. We used two different RNAi lines to knock them down everywhere with *TubP-Gal4*, initiating knockdown in 6-day old adult females. After two days, flies were fed DSS for two days, then repair was allowed for two days (6 days total knockdown). By analyzing muscle shape (aspect ratio), we found *trol*, *Ndg*, and *SPARC* were all required for basement membrane repair (Fig. S1A-F). The requirement for new basement membrane proteins indicates that damaged basement membranes are replaced with new matrix. These same proteins are required for basement membrane maintenance without externally applied damage, as knocking down *trol*, *Ndg,* or *SPARC* for 10 days in adults caused dysmorphic gut muscles (Fig. S1K-P), whereas knocking down non-matrix extracellular proteins *Gbp1* or *Gbp2*^19^ had no effect (Fig. S1G-J, Q-T). From this, we conclude that although basement membrane is replaced during normal homeostatic maintenance, it is replaced faster during repair.

### Collagen IV is upregulated in matrix mender cells following damage

Although basement membranes are maintained throughout the body, we expect damage-triggered repair to be a localized process. The Col4 in many *Drosophila* tissues originates from a distant organ, the fat body^20,21^, and we found that was true for the undamaged gut by expressing tissue-specific dsRNA against GFP in *vkg-GFP/vkg+* heterozygotes. *vkg-GFP/vkg+* gut basement membranes fluoresce green, but when dsRNA against GFP was expressed ubiquitously with *TubP-Gal4*, there was virtually no gut fluorescence, demonstrating the effectiveness of heterozygous knockdown (Fig S2 A-D). Importantly, the knockdown animal is functionally equivalent to a *vkg* heterozygote, viable and healthy^21^, so we do not expect to trigger any compensatory mechanisms. When GFP was knocked down specifically in the fat body with *c564-Gal4*, gut fluorescence was also extremely low, similar to ubiquitous knockdown (Fig. 1B,C and Fig. S2A-D; specificity of *c564-Gal4* shown in Fig. S2I,J), indicating most of the midgut Col4 originates from the fat body, a distant source. However, when these animals were fed DSS to damage the gut basement membrane, the gut fluorescence level increased significantly, indicating that Col4 was coming from another source besides the fat body during repair (Fig. 1C-E; see also Fig. SE-H). In cross-section, the DSS-induced Vkg-GFP was deposited between the gut epithelium and peristalsis muscles, suggesting a local source of Col4 specific for repair (Fig. 1F,G).

To identify a damage-specific local source of Col4, we used *Cg-Gal4*, a reporter of Collagen IV gene expression in which Gal4 is under control of a collagen enhancer that lies between the head-to-head promoters of the two collagen genes *Col4a1* and *vkg*^22^. In control guts, only a few cells in the posterior midgut (R4) faintly expressed *Cg>GFP,* but after 2 d of DSS damage the number and intensity of *Cg>GFP* cells in the gut increased substantially, from 1.8% to 5.7% of gut epithelial cells (Fig. 1H-M, Fig. S3A). Analyzing their numbers over time, we found that *Cg>GFP* cells increased after only one day of DSS feeding, peaked after the second day of DSS, and then subsided after DSS is removed. Interestingly, *Cg>GFP* expression was no longer observed 2 days after DSS removal, when the gut basement membrane was repaired (Fig. 1L). We dubbed these *Cg>GFP* cells “matrix menders” because they express Col4 in response to basement membrane damage but not after repair is complete. Analyzing other genes required for basement membrane repair, we found that reporters for *Pxn* and *Laminin* expression (*Pxn>GFP* and *LanB1>GFP*) were active in the gut without basement membrane damage; the number of *Pxn-*expressing cells doubled after damage, but the number of *LanB1*-expressing cells did not increase (Fig. S4).

### Matrix menders arise in response to basement membrane damage rather than epithelial damage

In addition to damaging basement membranes, DSS also damages cells^23^. To assess the damage profile that induces matrix menders, we tested two additional drugs known to damage the fly gut. Bleomycin is a DNA-damaging agent that targets enterocytes, causing tissue disorganization, cell loss, and a resulting increase in stem cell divisions^23^. After feeding bleomycin for 2 days, DNA damage was evident by anti- H2AvD staining and increased stem cell divisions were evident by anti-pH3 staining (Fig. S3B,C), yet the basement membranes remained undamaged as determined by muscle shape, and matrix mender cells were not evident (Fig. 1N-Q). Next we tested paraquat, which damages the fly gut by inducing reactive oxygen species and increasing stem cell divisions^24,25^. Feeding flies paraquat for 16 h, reported to maximize damage without causing lethality^24^, caused unexpected basement membrane damage, evident by altered muscle shape, and it significantly induced matrix menders (Fig. 1R- U), similar to the levels seen in response to DSS. These results indicate that matrix mender cells arise specifically to basement membrane damage, rather than epithelial damage.

### Matrix menders are a subset of enteroblasts

The cell types of the fly posterior midgut are well known (Fig. 2A) and can be identified with established cell markers. In DSS-fed guts, the Cg>GFP matrix menders did not express the intestinal stem cell marker Dl, but interestingly, about 75% of matrix menders were adjacent to Dl+ stem cells (174/236), suggesting that matrix menders are either enteroblasts or pre-enteroendocrine cells. Nearly all *Cg>GFP* labeled matrix menders were positive for the enteroblast marker *Su(H)GBE-lacZ*, indicating that matrix menders are enteroblasts (Fig. 2B,C). Not all enteroblasts are matrix menders, however, as only 43% of the *Su(H)GBE*-positive enteroblasts were also positive for *Cg>GFP* (Fig. 2C). In guts that were not damaged by DSS, *Cg-*expressing cells were still enteroblasts (81%) but fewer enteroblasts expressed *Cg* (21%) (Fig. 2C). Thus, some but not all enteroblasts are matrix menders. The number of enteroblasts is known to increase in response to DSS^23^, and we found that after 2 d of DSS, the number of enteroblasts increased from an average of ∼7/100 cells to ∼11/100 cells (Fig. 2D and Fig. S6A). Nearly all this increase in enteroblasts can be accounted for by the increase in collagen-expressing matrix mender cells (Fig. 1L).

**Figure 2.**
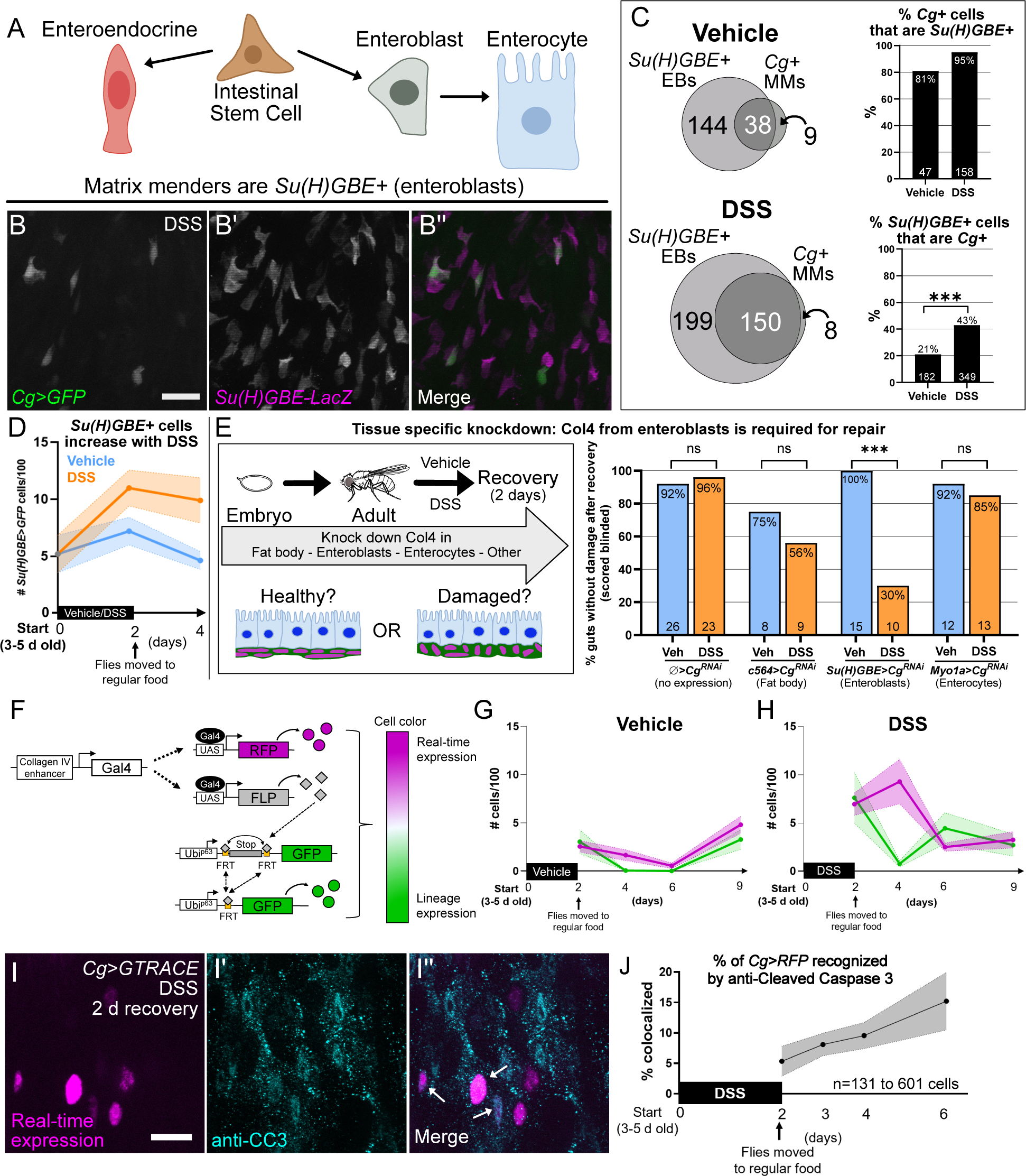
Matrix menders are a subset of enteroblasts that provide Col4 necessary for repair and are short-lived. **(A)** Lineage of gut epithelial cell types. **(B)** Matrix menders (*Cg>GFP*, green) expressed an enteroblast marker (*Su(H)GBE- lacZ,* magenta). Scale bar, 10 µm. **(C)** In both damaged and undamaged conditions, nearly all matrix menders were enteroblasts expressing *Su(H)GBE-lacZ*, but not all enteroblasts were matrix menders expressing *Cg>GFP*. After damage, more enteroblasts expressed *Cg>GFP*, evaluated by Fisher’s exact test. Number of cells scored is indicated. **(D)** Total enteroblasts increased in response to DSS, SEM indicated. Each dot represents 5-19 guts. Complete dataset in Fig. S6A. **(E)** Repair required Col4 expressed from enteroblasts. *Col4a1* was knocked down in different tissues, and adult gut muscle shape was compared in damaged-and-repaired guts to unchallenged guts. When Col4a1 was fully expressed (first two columns), nearly all guts had undamaged healthy basement membranes, both unchallenged and after repair. Col4 from the fat body was required for development and/or homeostasis, as guts appeared damaged even without challenge. Enteroblast Col4 was required specifically for repair. Col4 was not required from enterocytes. Numbers of guts evaluated is shown at column base. Guts were scored blinded to sample identity. Knock-down in other gut cell types and knockdown of *vkg* confirmed that Col4 is specifically required from enteroblasts for repair, see Fig. S5A-C. Significance evaluated by Fisher’s exact test. **(F-H)** Dynamics of matrix mender cells during and after collagen expression determined by G-TRACE. Overview (F) adapted from Evans et al., 2009. G, DSS increased the number of real-time *Cg*-expressing cells (RFP+, magenta); after repair, *Cg*-expressing RFP+ cells declined quickly, but most did not transition into lineage traced GFP+ cells (green). Cells expressing both RFP+ and GFP+ were counted only as RFP+ (magenta) to count each cell only once. Complete dataset in Fig. S6C,D. **I,J)** *Cg>GTRACE* real-time expressing cells (RFP+, magenta) underwent cell death, expressing Cleaved Caspase 3. Cell death of Cg-expressing cells was also observed without DSS, see Fig. S6G. Shading represents SEM in G,H,J.

### Collagen IV from enteroblasts is required for repair

We wanted to determine if the Col4 from matrix-mender enteroblasts was important for repair, but unfortunately knocking down Col4 with *Cg-Gal4* would target collagen in other tissues, such as the fat body. Instead, we knocked down *Col4a1* in specific cell types with different Gal4 drivers, including the enteroblast driver *Su(H)GBE- Gal4* (Fig. 2E). With collagen removed from each potential source, we compared gut morphology in undamaged guts to those that were DSS-damaged and allowed to repair, scoring samples blinded to identity or treatment. When *Col4a1* was knocked down in the fat body with *c564-Gal4*, many guts had dysmorphic muscles even without damage, and morphology was similar when treated with DSS and allowed to repair; these results indicate that Col4 from the fat body is important for gut basement membrane development or homeostasis as expected, but it is not specific for repair. We next knocked down Col4a1 in enteroblasts with *Su(H)GBE-Gal4,* which includes matrix menders; although *Su(H)GBE* is expressed in some larval tissues including wing and tracheae^26^, it is not expressed in known collagen source tissues (adult fat body, Malpighian tubules, or ovary, Fig. S5D-G). Gut muscle morphology was normal without DSS treatment, but repair was severely compromised, with 70% still damaged two days after removing DSS. To confirm the specificity of RNAi-mediated knockdown, we knocked down *vkg* (Col4a2) with *Su(H)GBE-Gal4*, and this also interfered specifically with repair and not development (Fig. S5A,B). These results indicate that Col4 from the matrix mender cells is required specifically for midgut basement membrane repair.

When *Col4a1* was knocked down with *Myo1a-Gal4* in enterocytes, the terminally differentiated cell type arising from enteroblasts, guts appeared like no-knockdown controls, normal both without damage and after repair; these results indicate that after differentiation, enterocytes do not supply Col4 for development, homeostasis, or repair. We also knocked down *Col4a1* in the intestinal stem cells and enteroendocrine cells (Fig. S5C), and morphology appeared similar without damage and after repair. Only the collagen from enteroblasts is specific for gut basement membrane repair.

### Most matrix menders are removed from the gut epithelium after repair

We were interested in what happened to the matrix mender cells after repair, when they were no longer evident. Did they turn off collagen expression, did they differentiate, or did they die? To address this question, we performed lineage tracing of the *Cg-Gal4* expressing matrix menders using the G-TRACE system^27^, in which cells actively expressing Col4 were labeled with *nlsRFP* and cells that previously expressed Col4 were permanently labeled with nlsGFP (Fig. 2F). We analyzed the number and fate of collagen-expressing cells in the gut, and as expected, the RFP+ cells behaved like the *Cg>GFP* matrix mender cells, except that their numbers peaked slightly higher and two days later (compare Fig. 2G,H to Fig. 1L). Although the difference is explained by nuclear RFP persisting longer than cytoplasmic GFP, it also indicates that matrix menders persist after they turn off collagen expression. After the peak, the number of RFP+ cells decreased rapidly. We expected that the RFP+ cells would transition into GFP+ lineage cells, because enteroblasts can remain dormant for weeks before differentiating into enterocytes in a healthy gut epithelium^28^. Contrary to our expectations, however, as RFP+ cells disappeared, the corresponding pulse of GFP+ cells was much smaller (Fig. 2H). These results indicate that most matrix menders are removed from the gut epithelium after repair is complete. Matrix menders are capable of differentiating into enterocytes, however, as the enterocyte-specific antibody Pdm1 stained about 15% (50/340) of GFP+ lineage-expressing cells after 2 days recovery but did not stain RFP+ collagen-expressing cells (Fig. S6F,G). Matrix menders never became enteroendocrine cells, as no G-TRACE labeled cells were labeled with anti- Prospero (Fig. S6E). To determine if some matrix menders were dying, we stained *Cg>G-TRACE* flies with anti-Cleaved Caspase 3 (CC3). In vehicle-treated guts, we found ∼9% of RFP+ matrix mender cells were positive for the cell death marker (Fig. S6H). In guts recovering from DSS damage, the percentage of dying matrix menders increased over time: at 0 d post DSS, 5% were labeled as dying, but by 4 d post DSS feeding (2 d recovery) 15% RFP+ cells were labeled as dying (Fig. 2I,J and Fig S6H). Although kinetic analysis is limited in fixed samples, it appears that most matrix menders do not differentiate into enterocytes and are short-lived, a property of enteroblasts generally after DSS treatment (Fig. S6B). Taken together, these results indicate that as repair is completed, matrix menders first turn off collagen, then a few differentiate while most die.

### Piezo is required for matrix menders to arise in response to damage

How does the gut sense damage to turn on Col4? Using a quantitative tensile strain assay, we previously demonstrated that DSS damage makes basement membranes less stiff (reduced elastic modulus)^14^, and we envision that most types of matrix damage would result in reduced stiffness (increased flexibility). Thus, we reasoned that basement membrane damage might be detected by a mechanosensitive mechanism such as the ion channel Piezo, known to be active in the fly gut^11,29,30^. To test Piezo’s role in basement membrane repair, we analyzed *Piezo^KO^* null flies, which are homozygous viable. After damaging with DSS, *Piezo^KO^* flies were unable to repair basement membranes (Fig. 3A-C), but unlike other matrix repair genes (Fig. S1), *Piezo* was not required for basement membrane maintenance (Fig. 3 D-F). Analysis of a second independent mutation, *Piezo^MI^*^04294^, confirmed its unusual repair-specific function (Fig. S7A-F). To determine if *Piezo* is required for detection of basement membrane damage, we quantified Cg-expressing matrix mender cells in *Piezo^KO^* versus control midguts after DSS damage. Piezo was required for matrix mender induction: in controls, 5.8% of epithelial cells were matrix menders, but in *Piezo^KO^* only 1.9% were and their *Cg* expression level was reduced (Fig. 3G-J). Thus, *Piezo* is required for matrix menders to arise in response to damage.

**Figure 3.**
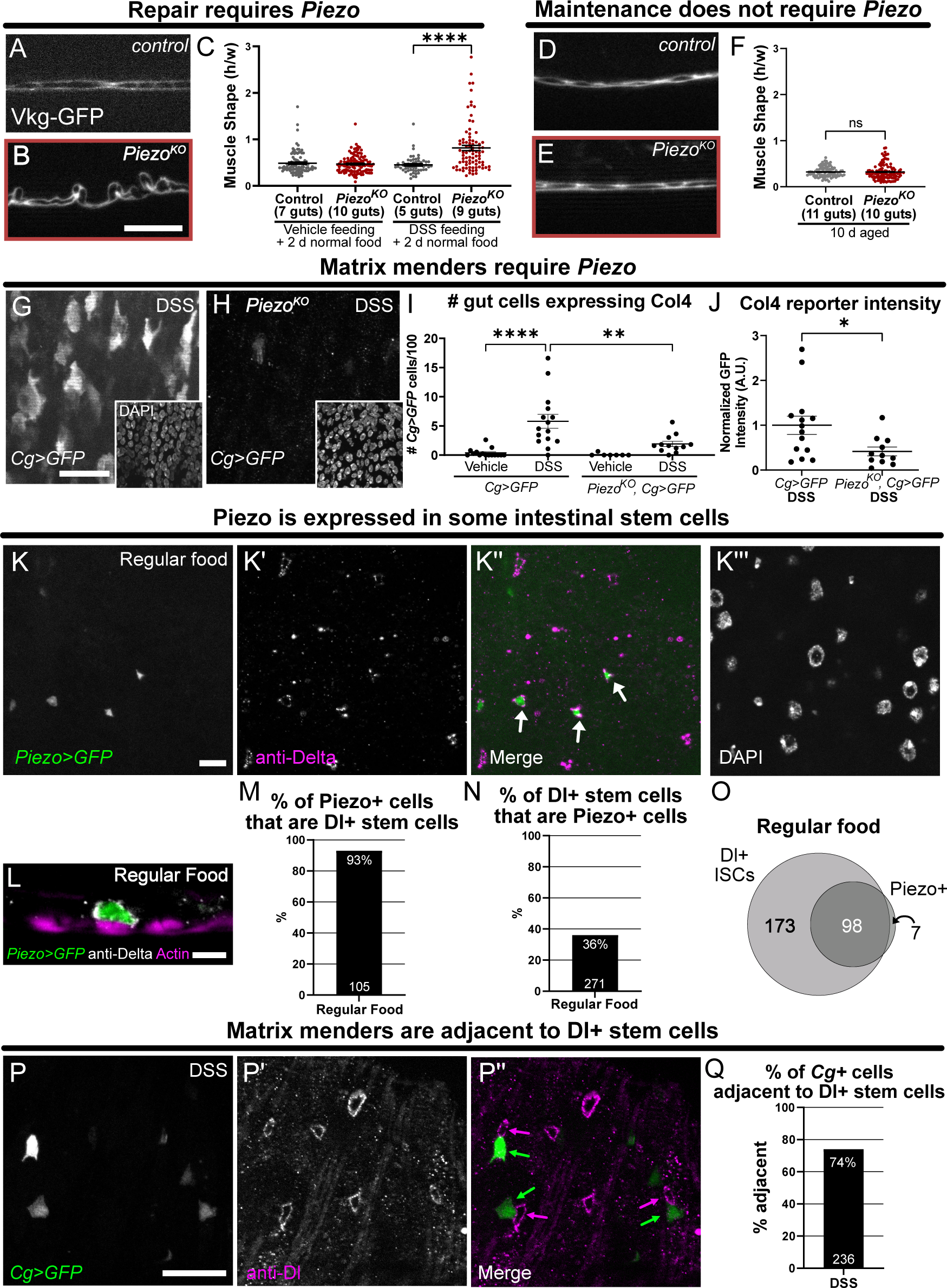
Piezo is required for basement membrane repair and for matrix menders to arise and is expressed in the intestinal stem cells. **(A-F)** *Piezo* was required for basement membrane repair but not maintenance in adult guts. See Fig. S7A-F for confirmation with another allele. Mean +/-SEM. Significance by *t-test*. Scale bar in B for A-E, 10 µm. **(G-J)** *Piezo* was required for *Cg>GFP* matrix mender cells to arise in response to DSS damage. Both the number and expression level of *Cg>GFP* cells were significantly reduced in *Piezo^KO^* flies. Each dot represents a gut (I,J). Mean +/-SEM. Significance by ANOVA (I) and *t*-test (J). Scale bar, 20 µm. **(K-O)** Piezo-transcriptional reporter (*Piezo^KI^-Gal4*, green) was expressed in Dl+ intestinal stem cells (magenta), flies fed normal food. Almost all Piezo+ cells were Delta+ intestinal stem cells, yet only 36% of Dl+ intestinal stem cells express Piezo. For Venn diagram, the circle size and amount of overlap are representative. Scale bars, 10 µm in K, 5 µm in L. **(P-Q)** *Cg>GFP* matrix mender cells (green) were adjacent to intestinal stem cells (anti- Delta, magenta). Scale bar, 10 µm. The number of cells analyzed is reported at column base for M,N,Q.

### Piezo is expressed in a subset of intestinal stem cells

For Piezo to monitor basement membrane damage, it must be present prior to damage. To localize Piezo, first we tried a functional GFP-Piezo fusion protein at the endogenous locus reported to be extremely dim in neurons^31^, but we were unable to detect GFP in the gut. Next, we turned to a Gal4-based *Piezo>GFP* transcriptional reporter, expressed outside the gut epithelium in tracheal cells that penetrate the gut muscles (Fig. S7G,H) and in small cells within the gut epithelium. On regular food, 93% of *Piezo>GFP* cells were positive for the intestinal stem cell marker Dl but only 36% of Dl+ stem cells expressed Piezo (Fig. 2K-O). Our results are similar to a previous report that *Piezo* is expressed in 40% of intestinal stem cells^11^. We found that stem cells are often adjacent to matrix menders (Fig. 3P-Q), suggesting that stem cell divisions give rise to matrix menders in a Piezo-dependent manner.

### Piezo detects decreased basement membrane stiffness to activate matrix menders

What aspect of basement membrane damage does Piezo detect? Piezo is a transmembrane channel activated by mechanical stretch to the plasma membrane^32^, and the plasma membranes of intestinal stem cells are in direct contact with the underlying basement membrane^33^. A damage-induced loss of tensile stiffness in the basement membrane is expected to put more tension on neighboring plasma membranes, resulting in membrane stretching and Piezo activation. Thus, we hypothesized that Piezo detects basement membrane damage via its loss of stiffness. As a test, we specifically reduced basement membrane stiffness in otherwise healthy and undamaged animals, by inhibiting the Col4 crosslinking enzyme Peroxidasin.

*Peroxidasin* mouse mutant tissue has reduced basement membrane stiffness (Young’s modulus), measured as increased deformation (stretch) in response to tension^18^. Using the same quantitative tensile strain assay, we recently reported that inhibiting Peroxidasin in wild-type adult *Drosophila* by feeding 100 µM of the suicide substrate phloroglucinol (PHG) for 5 days decreases basement membrane stiffness by about half^17^. The PHG-induced decrease in basement membrane stiffness was evident in the muscle aspect ratio, both at 100 µM and even more strongly at 5 mM, indicating a dose response (Fig. 7A-D). Excitingly, 100 µM PHG alone without other damage induced matrix menders (3.5% of gut epithelial cells), and 5 mM PHG induced even more matrix menders (5.3%), at a level similar to DSS induction (Fig. 4E,F,H, compare to Fig. 1L).

**Figure 4.**
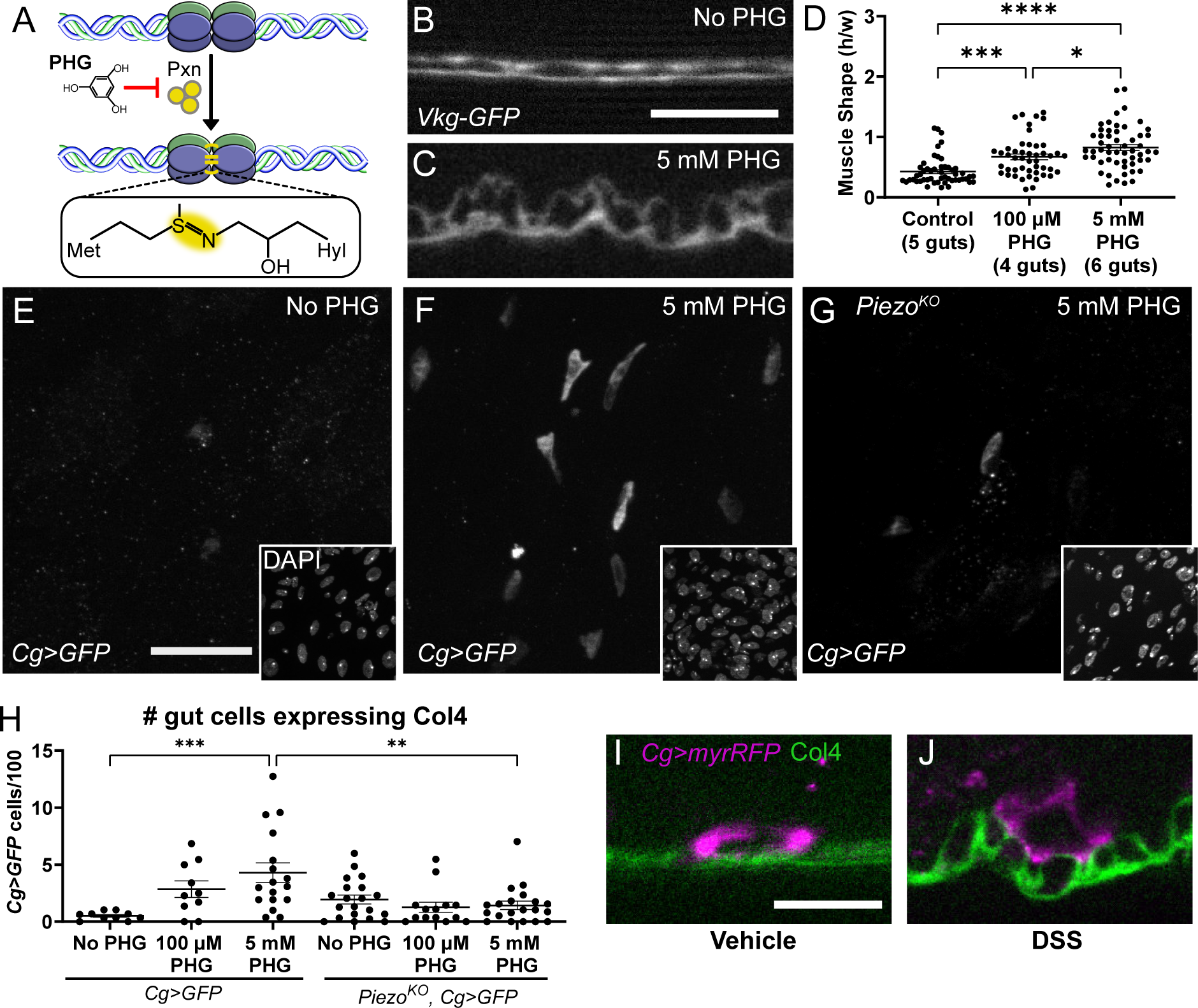
Piezo detects decreases in basement membrane stiffness to activate matrix menders. **(A)** PHG inhibits collagen IV crosslinking enzyme Pxn. **(B-D)** Feeding PHG reduced basement membrane stiffness in a dose-dependent manner, evidenced by muscle aspect ratio. Quantitative tensile stiffness measurements were reported in Peebles et. al., (2024). Significance evaluated by ANOVA. Mean +/- SEM. Scale bar, 10 µm. **(E-F)** Reducing collagen crosslinking induced matrix mender cells. Scale bar, 20 µm. **(G-H)** Piezo was required for matrix mender cells to arise in response to reduced collagen crosslinking. Each data point represents an individual gut. Mean +/-SEM. Significance by ANOVA. **(I-J)** Matrix mender cells have extensive contact with basement membrane, both with and without DSS (no recovery), positioning them well to repair it. Collagen is labeled with the triple-helix binding protein, CNA35 (green). Scale bar, 10 µm.

Thus, the loss of stiffness specifically activated matrix mender cells. Further, the mechanical induction of matrix menders required *Piezo*, as PHG-induced matrix menders were completely suppressed in the *Piezo^KO^* background (Fig. 4G,H). These results indicate that Piezo responds to the reduction in basement membrane stiffness to give rise to collagen-expressing matrix mender cells, positioned adjacent to the basement membrane (Fig. 4I,J) where they can effect repair.

## Discussion

Using the basement membrane around the *Drosophila* posterior midgut as an experimental model, we found that basement membrane damage gives rise to and is repaired by a distinct set of local epithelial cells, matrix mender cells, defined by their expression of the Collagen IV reporter *Cg-Gal4*. Matrix mender cells supply Col4 specifically required for repair: when Col4a1 or Vkg is knocked down in these cells, undamaged guts appear normal, but guts with basement membrane damage cannot repair. Most matrix menders are transient: after basement membrane repair is completed, matrix menders turn off collagen expression and most of them die. In contrast, Col4 is a long-lived molecule that is difficult to dispose of once deposited, and too much leads to fibrosis. Perhaps it is important to limit the duration of collagen- generating cells so they cannot cause fibrosis, thus making a long-lived molecule from a short-lived cell.

The matrix mender cells are a subset of enteroblasts, daughters of intestinal stem cells and progenitors of enterocytes. They are recognized by enteroblast marker *Su(H)GBE*, most sit next to an intestinal stem cell, and ∼15% differentiate into enterocytes. Supporting data for enteroblasts expressing Col4 come from mining published RNA sequencing data from single gut nuclei: transcripts for both Col4 subunits, *Cg25C* and *Vkg*, were identified almost exclusively in cells called progenitors of enterocytes (proECs), defined as a subset of enteroblasts; further, after DSS treatment, the number of proECs expressing the collagen transcripts doubled, results similar to our findings but using very different methods to identify collagen expression and cell identity^34,35^. Not all enteroblasts are matrix menders: in undamaged guts, only 21% of enteroblasts or 1.8% of epithelial cells express collagen, generally at low levels.

After basement membrane damage, the numbers increase: intestinal stem cell divisions generate more total enteroblasts, and a greater percentage of them express collagen (43%), with the result that about 6% of total epithelial cells express collagen, generally at higher levels.

How is matrix damage detected to give rise to matrix menders? Matrix menders arise in response to three independent challenges, DSS, paraquat, and PHG. All three damage the basement membrane as assayed by gut morphology; in contrast, matrix menders do not arise in response to the DNA damaging agent bleomycin, which does not appear to damage the basement membrane although it damages the epithelial layer and initiates stem-cell mediated repair^23,36^. Although DSS causes nonspecific damage to both cells and matrix in the gut epithelium, PHG is highly specific, reducing the number of head-to-head Col4 covalent sulfilimine crosslinks^16,37^ and decreasing basement membrane stiffness in a tensile strain assay, meaning that the basement membrane will stretch more (increased strain) in response to a given pulling force or tension^14,17^. The gut is a mechanically active environment, and increased basement membrane stretching would cause increased stretching of neighboring plasma membranes in the gut epithelial monolayer, activating mechanical stretch receptors.

Indeed, we found that the mechanosensitive transmembrane ion channel Piezo is required specifically for basement membrane repair. Although many genes are required for basement membrane repair, including those encoding basement membrane components Nidogen, Perlecan, Laminin, Collagen IV, Peroxidasin, and the Collagen chaperone protein SPARC, these genes are not specific to repair, as they are also required for basement membrane maintenance. In contrast, the mechanosensitive ion channel Piezo is required for repair only, not maintenance, confirmed with two independent mutations. Our data strongly support a model that Piezo detects reductions in basement membrane stiffness and signals for the generation of matrix menders.

Interestingly, even without induced damage, a few cells express collagen at low levels in the midgut epithelium. We expect that low levels of basement membrane damage are intrinsic to gut physiology given the proximity of the gut to an unpredictable external environment and the mechanical wear and tear of peristalsis, and this low level of damage would induce a weak matrix-mender response. Our various control data show that in undamaged conditions the number of matrix menders is somewhat variable, consistent with stochastic low levels of matrix damage.

### Simple model of basement membrane repair and its limitations

We and others found Piezo expression in intestinal stem cells, so our data are consistent with a simple stem-cell based model of basement membrane repair: intestinal stem cells directly monitor basement membrane stiffness, with matrix damage allowing greater cell stretching, activating Piezo. As a cation channel, Piezo activation increases intracellular calcium levels, and sustained cytoplasmic calcium is known to be sufficient to induce intestinal stem cell proliferation^38,39^. Thus, stem cells give rise to matrix menders that express and secrete Col4 locally, restoring the structure of the basement membrane. Their job complete, these cells turn off collagen expression and most of them die. Although our data support this attractive model, it remains unclear where Piezo detects basement membrane stiffness. Piezo is expressed in Dl+ intestinal stem cells, but its function there has not been demonstrated as we have not had success with Piezo conditional knockdown (i.e. ubiquitous knockdown does not reproduce the mutant phenotype). Nevertheless, our data strongly support a two-part model: (1) mechanical surveillance of basement membrane stiffness by Piezo, which activates (2) repair mediated by local epithelial cells. Indeed, very recently it was shown that stiffness regulates mouse intestinal stem cells via Piezo^40^, and this may represent part of the mechanism we show here. Our basement membrane repair model is likely to be generalizable to other tissues and animals, as Piezo and basement membrane components are all highly conserved as is the tissue architecture of basement membranes in close contact with epithelial cells.

## Lead contact

Further information and requests for resources and reagents should be directed to and will be fulfilled by the lead contact, Andrea Page-McCaw

## Materials availability

Any requests for resources and reagents should be directed to the lead contact.

## Data and code availability

All original code is publicly available: https://github.com/mshutson/cell-shape-analysis

## Acknowledgements

We thank K. Elkie Peebles, Kimberly LaFever, Jordyn Sanner Barr, Leah Caplan, and Junmin Hua for technical help, Sergei Boudko and Patrick Page-McCaw for discussions, and Barry Denholm, Bruce Edgar, Yuh Nung Jan, BDSC, VDRC, and Kyoto DSC for fly stocks. CNA35 recombinant monomer protein was a gift from Sergei Boudko, and anti-Pdm1 was a gift from Yang Xiaohang. Work was supported by NIGMS (R01GM137595 to A.P-M., and F31GM148021 to A.S.)

## Author Contributions

AS: Conceptualization, methodology, validation, formal analysis, investigation, resources, writing and editing, visualization. MSH: Methodology, software. APM: Conceptualization, methodology, writing and editing, visualization, supervision, project administration.

## Supplemental Information

Document S1. Figures S1-S7

## Materials and Methods

### Key resources table

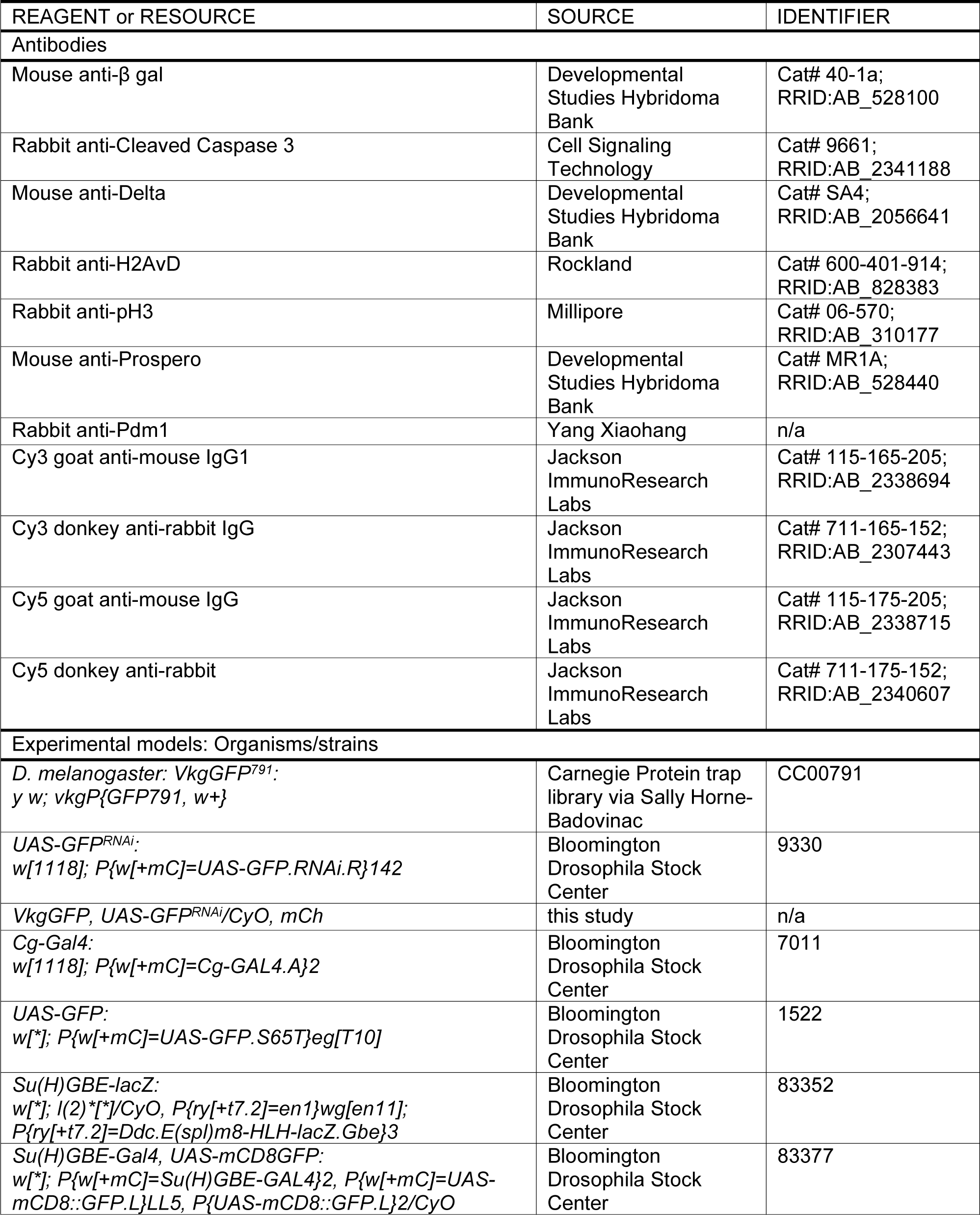
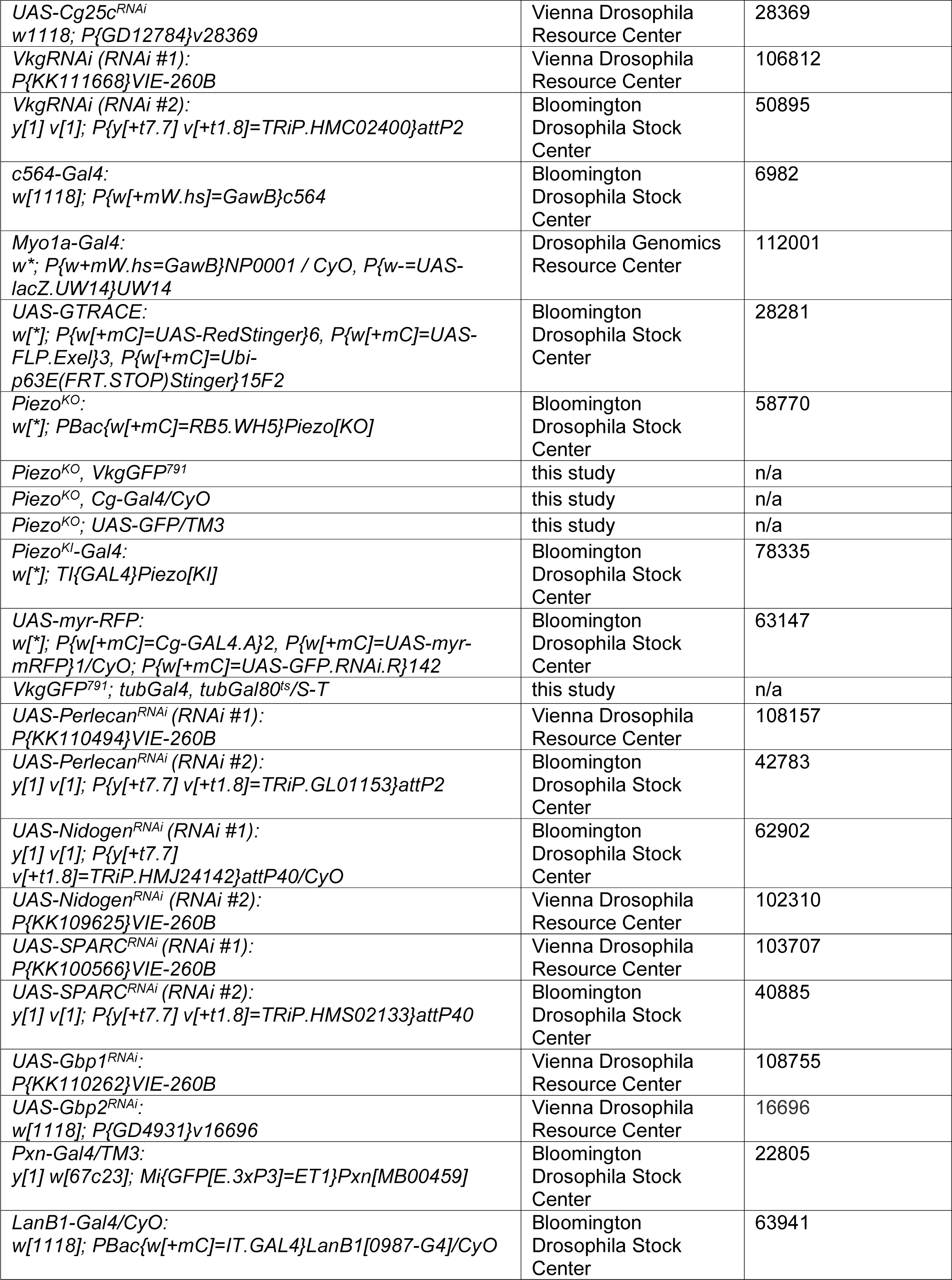
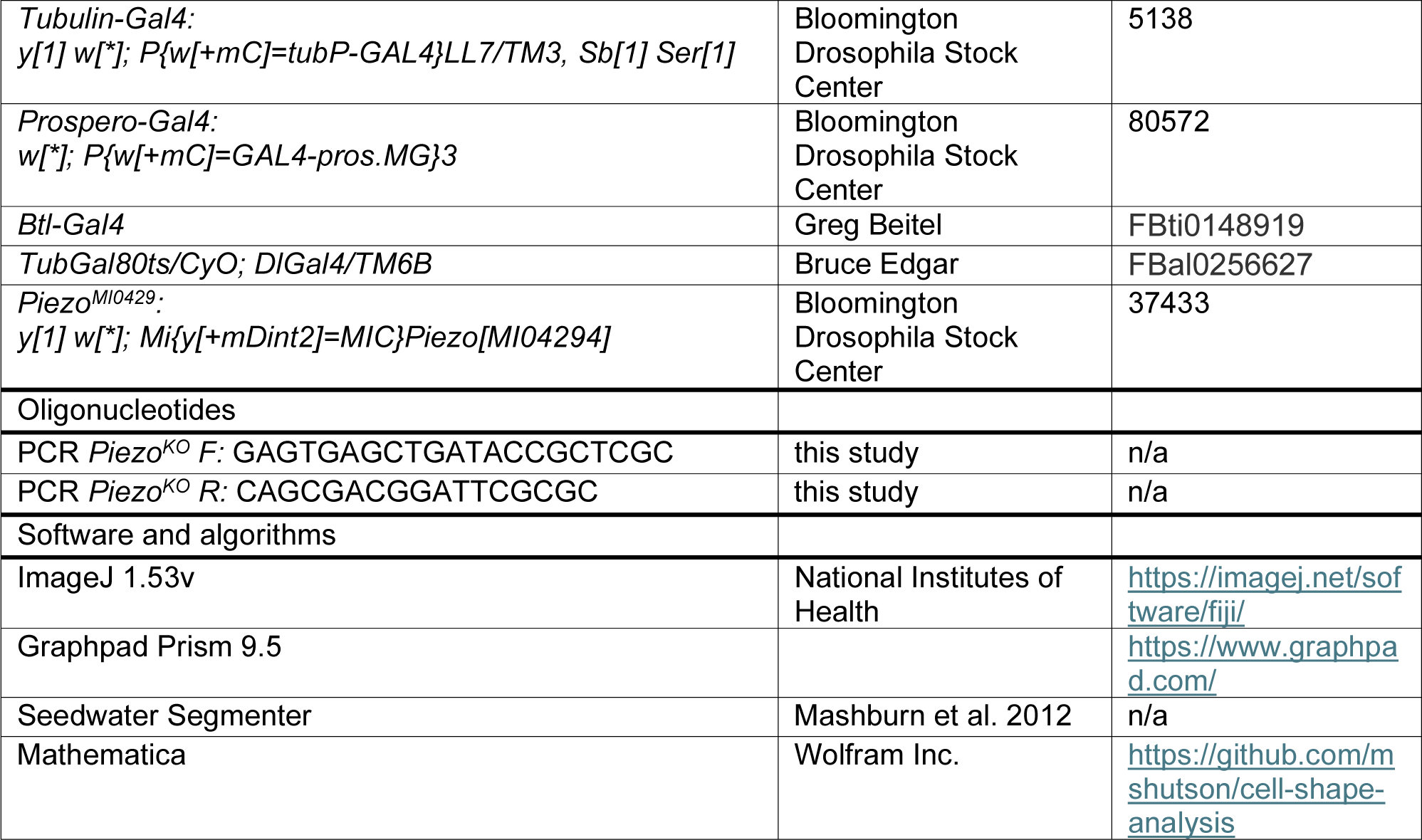

### Fly husbandry

Flies were maintained on cornmeal-molasses food at 25 °C unless otherwise specified. Adult-onset RNAi-based gene knockdown was done under *Gal80^ts^* control, with flies maintained at 18 °C until they eclosed. Female flies were then aged at 18 °C until they were 6-day adults (mated) before moving them to 29 °C to activate Gal4. Each experiment contains data from at least two independent crosses. For the list of fly stocks, please refer to Key Resource Table.

### Confirming *Piezo^KO^* using PCR

Because *Piezo^KO^* flies have no obvious phenotype, we verified the presence of the deletion by PCR in the starting stock and in recombinants. Primers were designed and optimized against the Piggybac insertion PBac{WH}CG8486-f02291 in *Piezo^KO^* animals to confirm the loss of Piezo using the following primers – F: GAGTGAGCTGATACCGCTCGC, R: CAGCGACGGATTCGCGC. OneTaq2x Master Mix with standard buffer was used. The following PCR conditions were used – Initial Denaturation: 94 °C, 30 sec; denaturation: 94 °C, 30 sec; annealing: 65 °C, 60 sec; extension: 68 °C, 45 sec (30x cycles); final extension: 68 °C, 5 minutes.

### DSS feeding

All DSS feeding experiments used mated female flies aged 3-5 d after eclosion before feeding DSS. Flies utilizing the *Gal80^ts^* system were moved to 29 °C for 2 days to ensure the Gal4 was active prior to starting DSS feeding treatments. Flies not utilizing the Gal80^ts^ system were aged 3-5 days at 25 °C prior to starting DSS feeding treatments. DSS was administered as described in Amcheslavsky et al. (2009) and Howard et al. (2019): a 2.5 cm x 3 cm piece of chromatography paper (Whatman 3030- 861, Grade 3 MM CHR) was placed in an empty vial. 500 µl of a 5% sucrose solution with or without 3% 36-50 kDa DSS (dextran sulfate sodium salt colitis grade, MP Biomedicals, CAS number 9011-18-1, 36,000 – 50,000 MW) was added to the paper in the vial. Female flies were anesthetized and carefully placed in the vial. Flies were added to new vials with fresh media each day for 2 days. For recovery experiments, flies were transferred to a vial containing standard cornmeal-molasses food for 2 days.

For imaging in 3 dimensions (Z-stacks), flies on standard cornmeal-molasses food were fed 5% sucrose for 4 hours before dissection to reduce gut autofluorescence from the food.

### Phloroglucinol feeding

PHG was fed to flies by dissolving it in food prepared without any cornmeal, freshly made with or without 100 µM or 5 mM Phloroglucinol (PHG) (Sigma-Aldrich, CAS number 108-73-6) and solidified in a vial overnight. 20 aged female flies and 5 male flies were added to the vials with or without PHG. Flies were kept at 25 °C for 5 days. Prior to dissection, flies were fed 5% sucrose for 4 hours before dissection to reduce gut autofluorescence from the food.

### Bleomycin feeding

Bleomycin (Sigma, 9041-93-4) was dissolved in water at a final concentration of 25 µg/mL and 5% sucrose, or 5% sucrose as a control. 500 µl of bleomycin or control solution were added to a 2.5 cm x 3 cm piece of chromatography paper at the bottom of an empty food vial. Flies were fed bleomycin for 2 days, changing the filter paper and vial daily.

### Paraquat feeding

A 10 mM paraquat (Sigma, 75365-73-0) solution was made in water with 5% sucrose, or 5% sucrose alone as the control. Flies were starved in an empty vial for 4 hours before transferring them to a vial with a 2.5 cm x 3 cm piece of chromatography paper soaked with either 500 µl of paraquat or control solution. Flies were fed for 16 hours prior to dissection.

### Gut dissections and preparations

Adult females were placed in cold Grace’s media and pinched between the abdomen and thorax using sharp #5 dissecting forceps (Dumont). The abdomen was separated from the thorax, and the integument surrounding the abdomen was peeled away, exposing the gut and Malpighian tubules, being careful not to pinch or stretch the gut. The Grace’s media was removed using a Pasteur pipette, followed by one quick wash with 1x PBS (137 mM NaCl, 2.7 mM KCl, 10 mM Na2HPO4, and 1.8 mM KH2PO4). The guts were fixed with 8% paraformaldehyde (Ted Pella, Paraformaldehyde 16%, Product #18505) in PBS for 60 minutes at room temperature, followed by 3 quick washes in PBS, and 4 x 15 min washes with PBS or PBT.

For immunofluorescence experiments, blocking buffer was added to fixed guts for 2 hours at room temperature or overnight at 4 °C then incubated in primary antibody and 0.2 µM SiRActin (Spirochrome Cat# SC001) in PBS or PBT overnight at 4 °C, followed by 4 x 30 min washes with 1x PBS. Secondary antibodies were added for 2 hours at room temperature, followed by 4 x 30 min washes. Guts were then mounted in DAPI-containing mounting media (Vectashield, Vector Laboratories, H1200). Antibodies used were mouse anti-β gal (1:50), rabbit anti-Cleaved Caspase 3 (1:200), mouse anti-Delta (1:50), rabbit anti-H2AvD (1:1000), rabbit anti-pH3 (1:1000), mouse anti-Prospero (1:50), and rabbit anti-Pdm1 (1:1000). The following secondary antibodies were used at 1:200 Cy3 goat anti-mouse IgG1, Cy3 donkey anti-rabbit IgG, Cy5 goat anti-mouse IgG, and Cy5 donkey anti-rabbit.

### CNA35 staining

Immediately after gut dissections, CNA35 recombinant monomer protein labeled with Alexa488 (1 mg/ml in PBS) was diluted to 1:100 in PBS and added to guts for 2 hours, followed by 1 quick wash in PBS and fixed for 1 hour with 8% PFA.

### Light Microscopy

Single optical sections of the posterior midgut (R4) were imaged using a Zeiss Apotome mounted to an Axio Image M2 with a 63x/1.4 oil Plan-Apochromat objective.

Z-stacks were acquired using a 40x/1.3 oil Plan-Apochromat objective at a 0.75 µm step size taken through the top half of the gut tube (lumen through the top of the muscle layer). Images were taken using an AxioCam MRm camera (Zeiss), X-Cite 120Q light source (Excelitas Technologies), and AxioVision 4.8 software. FIJI ImageJ (version 1.53v, National Institutes of Health) 16-bit, ZVI files were used for analysis and figures for images.

For Fig 1C,D and Fig S2A-H, the fluorescence intensity of the basement membrane was determined by taking single optical slices *en face* through the middle of the circumferential muscles to ensure the same plane of the basement membrane was imaged, and the exposures were kept the same. The fluorescence intensity of the basement membrane was measured in ImageJ using the “box” tool to measure the average intensity of 3 representative regions of the gut basement membrane. Similarly, the background intensity was determined by subtracting the average of 3 boxes outside of the gut region. An AVOVA test was performed (GraphPad Prism 9.5).

Fluorescence intensity measurements of gut cells to be compared were acquired at the same exposure from at least 3 experiments done on different days. Maximum intensity projections were created using FIJI ImageJ and the line tool was used to draw through the center of the 3 brightest nuclei in the GFP channel and the fluorescence intensity was measured and averaged. Background fluorescence was subtracted by taking the average intensity of 3 lines within the gut region that did not overlap with GFP nuclei. An unpaired *t-test* was performed.

### Measurement of muscle shape

Using single-slice cross section images of the gut basement membrane, individual muscle cells were identified and segmented using the software Seedwater Segmenter (Mashburn et al. 2012). Custom code in Mathematica (Wolfram Inc., Champaign, IL) was then used to analyze the segmented images and determine the aspect ratio of each muscle cell (code available at https://github.com/mshutson/cell-shape-analysis). Specifically, to quantify the aspect ratio of gut muscle cells, we first fit a 4th-order polynomial to all the segmented cells to estimate a tangent line along the gut’s long axis. We then calculated the two-dimensional moment-of-inertia tensor, J, for each cell relative to the local tangent line, and calculated each cell’s aspect ratio as Sqrt[J11/J22]. This aspect ratio is thus the cell’s height normal to the gut’s long axis divided by the cell’s length along the gut’s long axis.

### Quantifying number of cells

GFP+ cells were manually quantified by creating a maximum intensity projection of the Z-stack images of the top half of the gut and confirmed by the presence of a nucleus. The total number of nuclei in the posterior midgut was quantified by creating a maximum intensity projection and setting a threshold to select the DAPI-stained nuclei.

Images were “despeckled,” and a watershed was applied to the image. Using the ImageJ “analyze particle” feature, the number of particles larger than 5 µm^2^ was quantified. The areas where nuclei were fused together or unable to be accurately quantified by ImageJ were manually quantified. The total number of GFP+ cells captured in each image was divided by the total number of nuclei in the image and multiplied by 100 to report the #GFP cells/100.

### Statistical reporting

All statistical analysis was performed using GraphPad Prism 9.5. ns indicates P > 0.05, * indicates P ≤ 0.05, ** indicates P ≤ 0.01, *** indicates P ≤ 0.001, **** indicates P ≤ 0.0001.

---

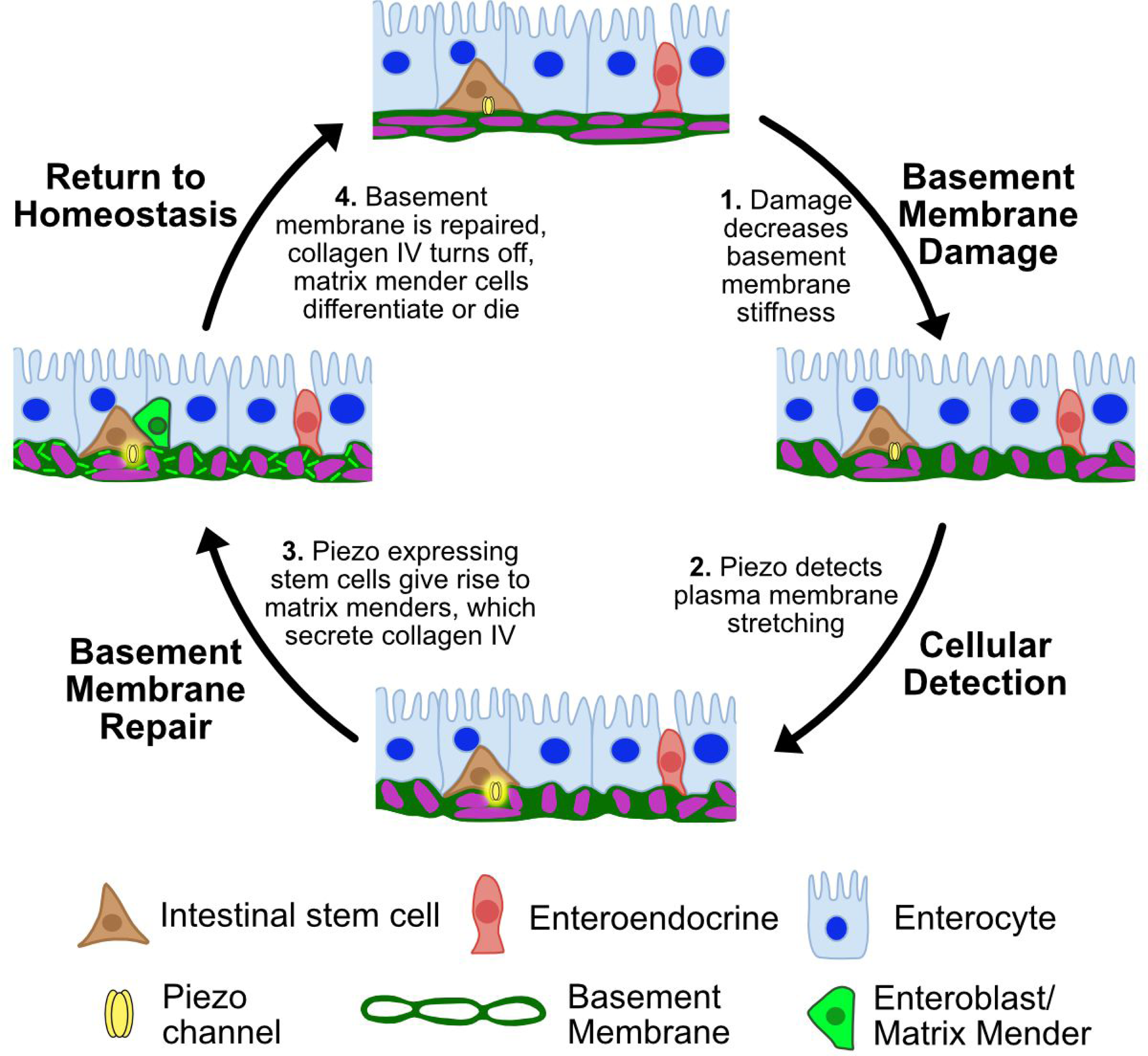

