## Supplemental Figure 1 for "Piezo-dependent surveillance of matrix stiffness generates transient cells that repair the basement membrane"

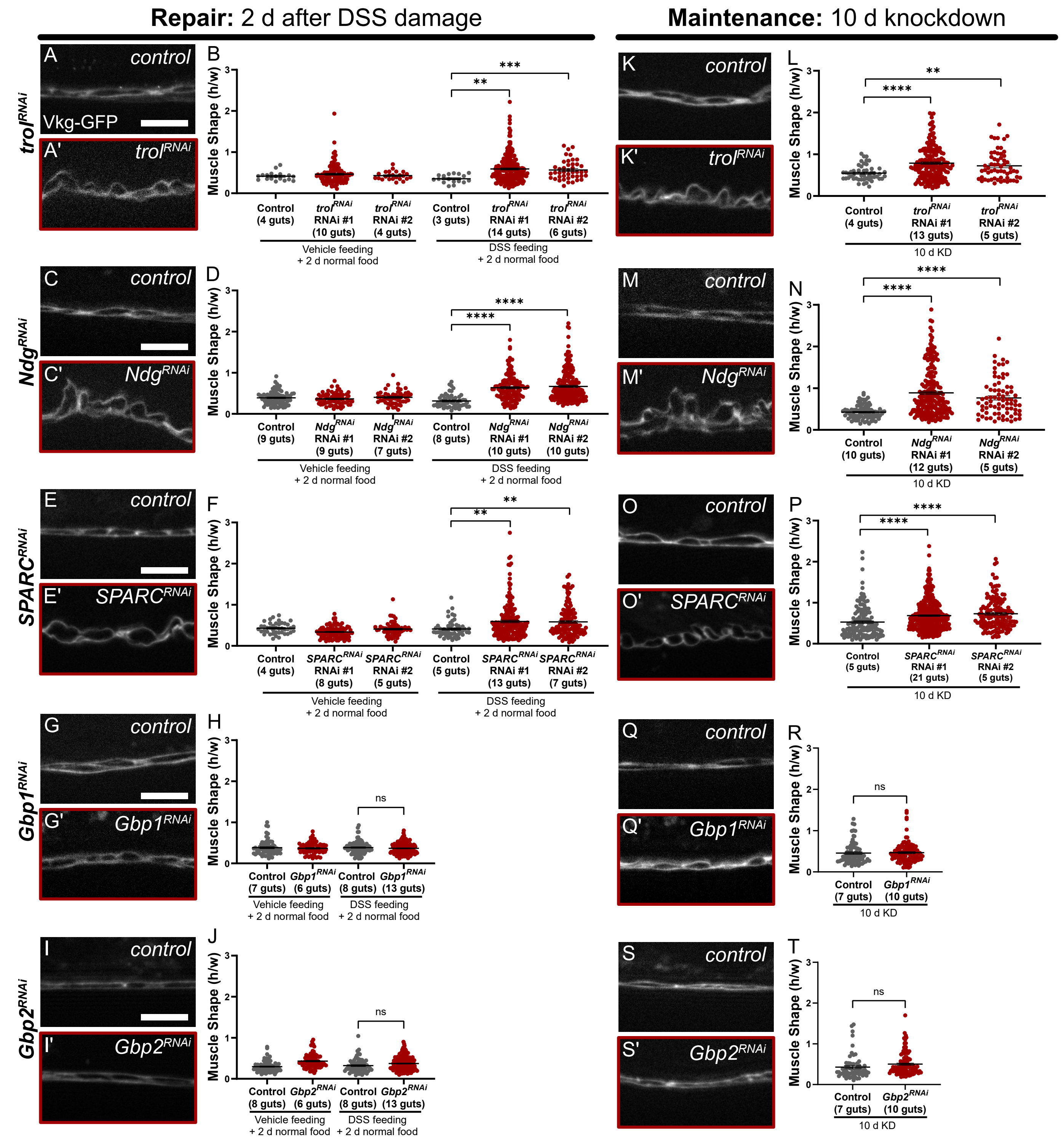


**Fig. S1. *Trol*, *Nidogen*, and *SPARC*, but not *Gbp1* and *Gbp2*, are required for basement membrane repair and maintenance, related to Figure 1.**

**(A-F)** *Trol*, *Nidogen,* and *SPARC* are required for repair (A-F) and maintenance (K-P) of basement membrane in adult guts. Basement membrane was visualized using Vkg-GFP, and damage was quantified as the muscle shape aspect ratio (height/width), each muscle shown as a dot. The number of guts analyzed is indicated. Mean +/- SEM, significance evaluated by *t*-test against control group, ** indicates P ≤ 0.01, **** indicates P ≤ 0.0001. Scale bar, 10 µm.

**(G-J)** *Gbp1* and *Gbp2* are not required for basement membrane repair after DSS. Mean +/- SEM, significance evaluated by *t*-test.

**(Q-T)** *Gbp1* and *Gbp2* are not required for basement membrane maintenance.
