## Supplemental Figure 2 for "Piezo-dependent surveillance of matrix stiffness generates transient cells that repair the basement membrane"

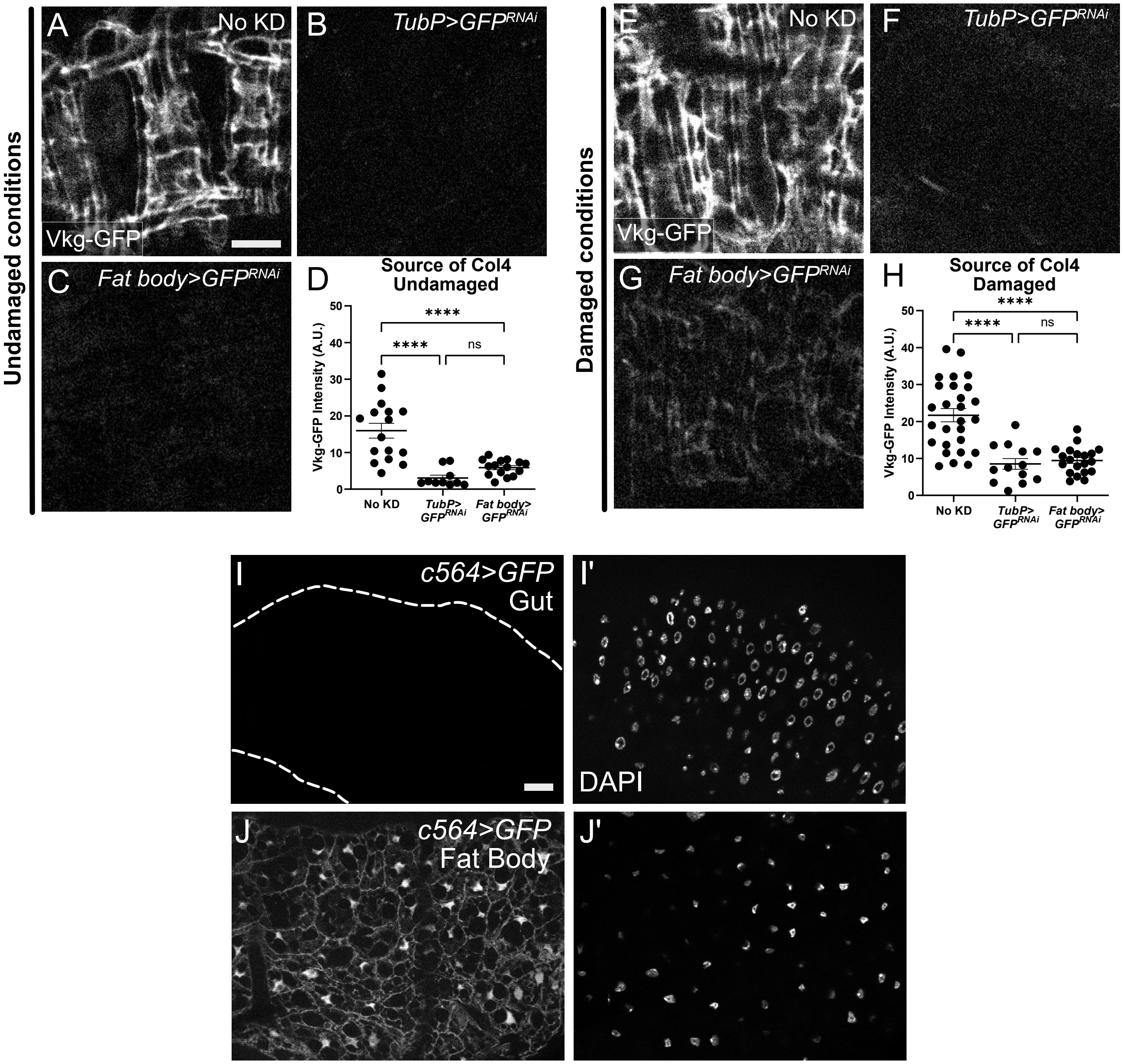


**Fig. S2. The fat body is the main source of Col4 in the adult gut basement membrane. *c564-Gal4* is specific to the fat body and is not expressed in the gut, related to Figure 1.**

**(A-D)** In undamaged conditions, midgut Col4 comes from the fat body. Surface view of gut basement membrane in *vkg-GFP/vkg+* controls (A). Ubiquitous knockdown of GFP eliminated Vkg-GFP fluorescence in the gut. Col4 expression from from the untagged wild-type allele was not affected by this strategy, alleviating the need for any potential compensatory mechanisms (B). Similar fluorescence loss was observed with fat-body specific GFP knockdown, using *c*564-Gal4 (C,D). Scale bar, 10 μm.

**(E-H)** In DSS-damaged conditions, similar fluorescence loss was observed with fat-body specific GFP knockdown, using *c564-Gal4*, suggesting the fat body is still a major contributor to the midgut Col4. In D and H, each dot represents a gut, mean +/- SEM, significance by ANOVA.

**(I, I')** *c564-Gal4* is not expressed in the posterior midgut (outlined), demonstrated by lack of expression of *UAS-GFP*, DAPI shown in I'. Scale bar, 20 μm.

**(J, J')** *c564-Gal4* is expressed specifically in the fat body, DAPI shown in J'.
