## Supplemental Figure 3 for "Piezo-dependent surveillance of matrix stiffness generates transient cells that repair the basement membrane"

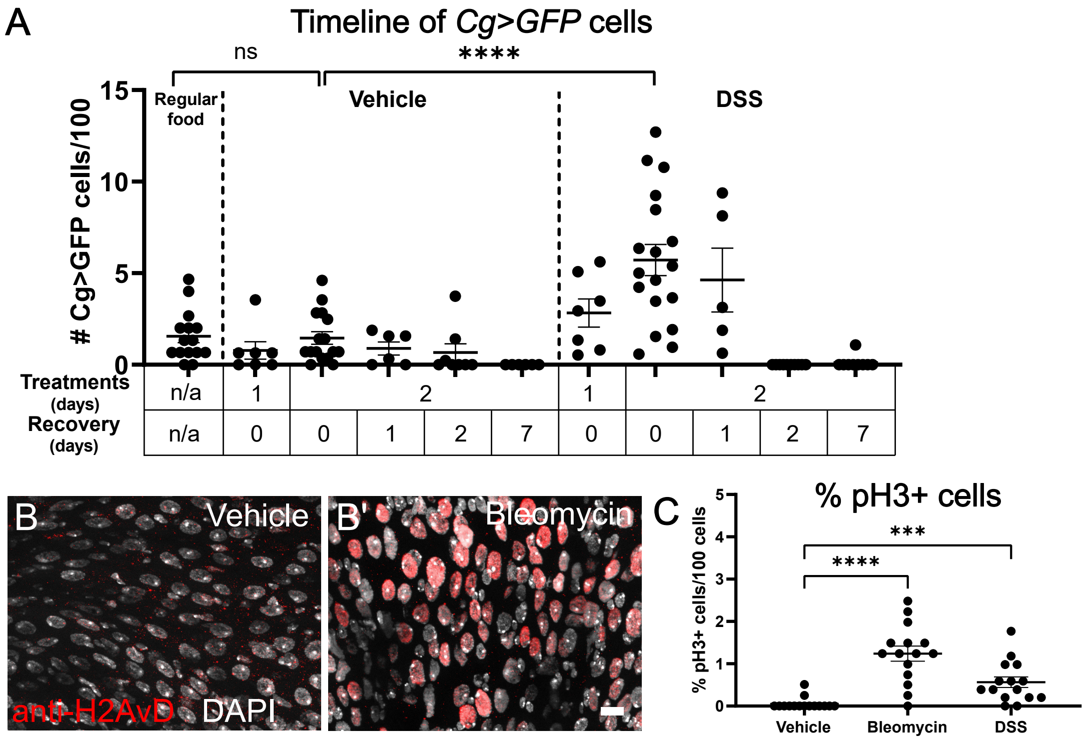


**Fig. S3. Supporting data for matrix mender dynamics and cells with DNA damage. Bleomycin and DSS treatments results in an increased number of dividing cells, related to Figure 1.**

**(A)** Individual data points from *Cg>GFP* timeline in Fig. 1L. Mean and SEM are indicated. Significance evaluated by ANOVA.

**(B)** After bleomycin treatment, gut epithelial cells were recognized by the DNA-damage specific histone antibody, anti-H2AvD, accompanying Fig 1N-Q. Scale bar, 10 μm.

**(C)** Feeding bleomycin or DSS for 2 days results in an increase in the pH3+ cells. A *t*-test between vehicle and bleomycin treatments or vehicle and DSS treatments were performed to determine significance.
