## Supplemental Figure 4 for "Piezo-dependent surveillance of matrix stiffness generates transient cells that repair the basement membrane"

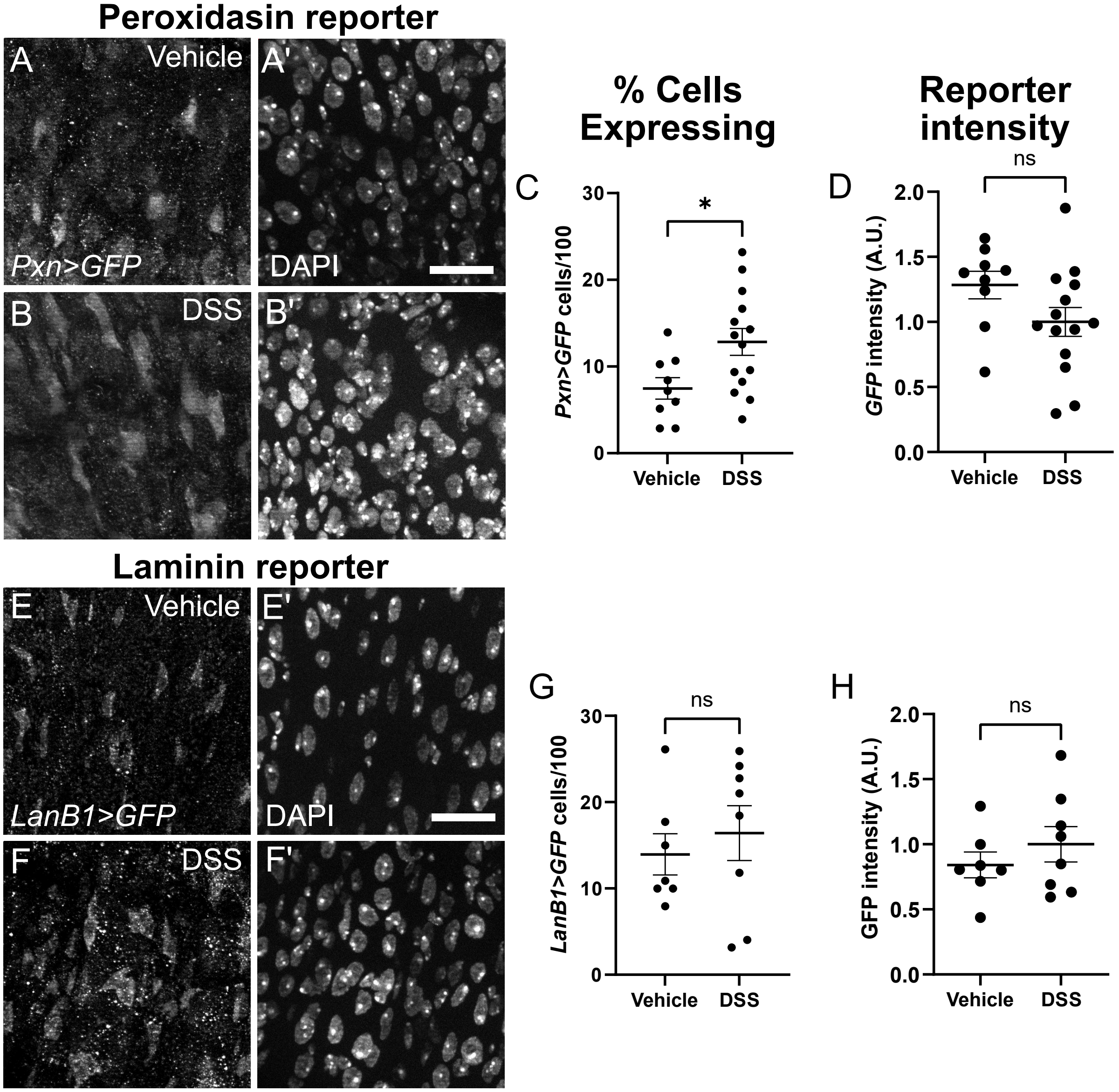


**Fig. S4.** ***Pxn>GFP*** **and** ***LamininB1>GFP*** **are expressed in the posterior midgut without DSS damage, and Pxn expression increases in response to DSS, related to Figure 1.**

**(A,B)** *Pxn* transcriptional reporter, *Pxn-Gal4,* was expressed even without damage (vehicle) in some posterior midgut cells. In response to DSS-induced basement membrane damage, the number of *Pxn*-expressing cells increased. Scale bar, 20 μm.

**(C,D)** Quantification shows the percentage of *Pxn>GFP* cells about doubled after DSS damage (C). However, the intensity of the *Pxn>GFP* transcriptional reporter did not change significantly within an expressing cell after DSS damage (D).

**(E,F)** *Laminin B1* transcriptional reporter, *LanB1-Gal4,* was expressed even without damage (vehicle) in some posterior midgut cells, and its expression was not increased in response to DSS-induced basement membrane damage. Scale bar, 20 μm.

**(G,H)** Quantification of number of *LanB1>GFP* cells (G) and intensity of the GFP transcriptional reporter (H) with or without DSS.
