## Supplemental Figure 5 for "Piezo-dependent surveillance of matrix stiffness generates transient cells that repair the basement membrane"

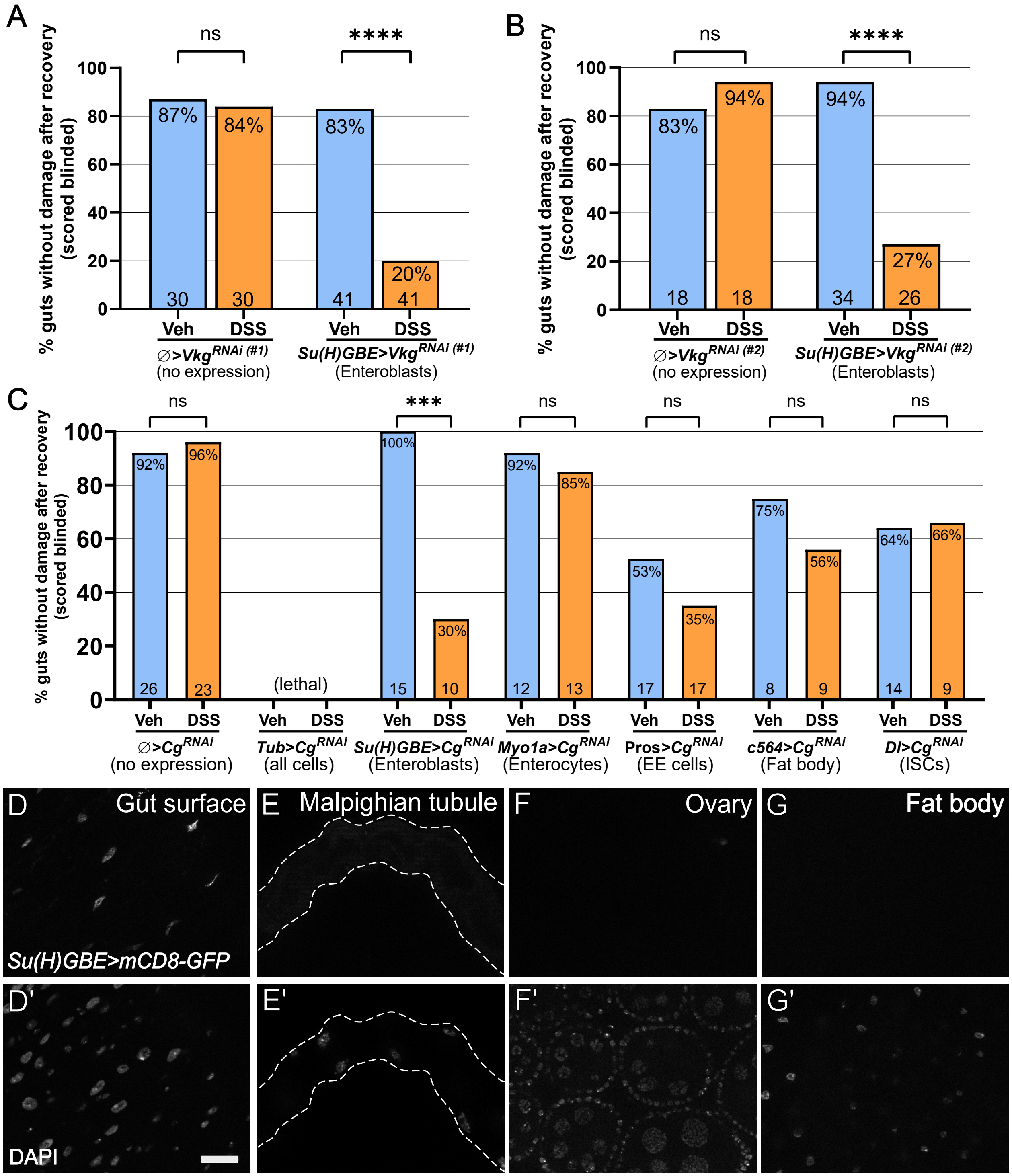


**Fig. S5. Collagen IV from Su(H)GBE-expressing enteroblasts is required for basement membrane repair, and Su(H)GBE is not highly expressed in the Malpighian tubules, ovaries, or fat body, related to Figure 2.**

**A,B)** Knock down of v*kg* (2 RNAi lines) in different cell types and tissues. The percentage of guts that had normal morphology after vehicle treatment and those that had normal morphology after DSS damage and repair is shown. *vkg* from enteroblasts expressing *Su(H)GBE* is required for repair. Morphology was scored blinded to sample identity. Significance was determined using a Fisher's exact test.

**C)** Complete data set of Col4a1 ($Cg^{RNAi}$) knock down in different cell types and tissues. The percentage of guts without damaged was determined as described in A,B. Col4a1 from enteroblasts expressing *Su(H)GBE* is required for repair.

**D-G)** *Su(H)GBE>mCD8-GFP* is expressed in the gut, but not the Malpighian tubules, ovaries, or fat body (D-G). Corresponding DAPI images are shown (D'-G'). Scale bar, 20 μm.
