## Supplemental Figure 6 for "Piezo-dependent surveillance of matrix stiffness generates transient cells that repair the basement membrane"

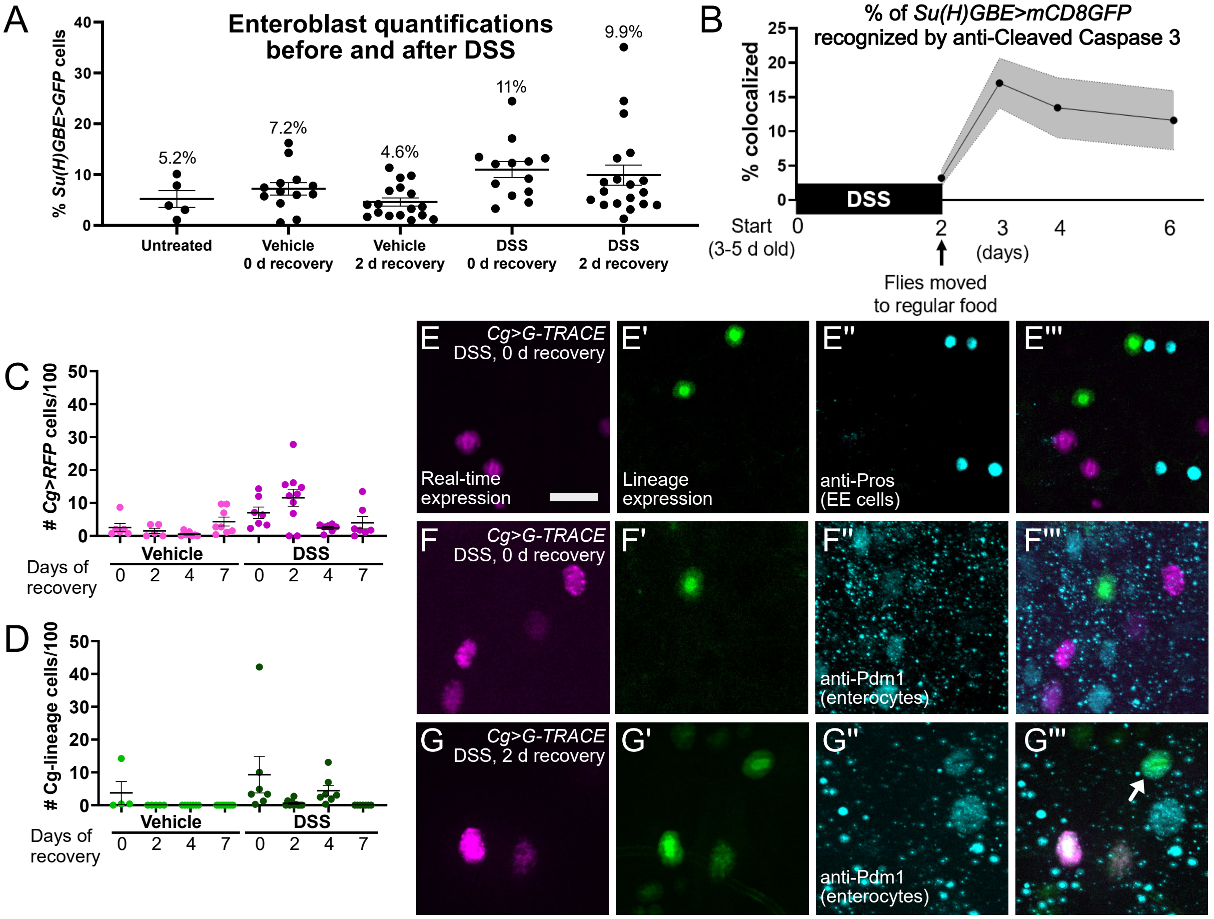


H

| | Vehicle | | | | DSS | | | |
|---|---|---|---|---|---|---|---|---|
| Days of regular food | 0 | 1 | 2 | 4 | 0 | 1 | 2 | 4 |
| % *Cg>RFP* cells that are CC3 + | **8%** (n=144) | **14%** (n=131) | **6%** (n=301) | **12%** (n=140) | **5%** (n=353) | **8%** (n=172) | **10%** (n=601) | **15%** (n=171) |
| % Lineage cells that are CC3 + | **14%** (n=58) | **34%** (n=82) | **26%** (n=98) | **4%** (n=56) | **14%** (n=171) | **5%** (n=157) | **5%** (n=227) | **6%** (n=63) |

**Fig. S6. The percent cells expressing the enteroblast marker *Su(H)GBE* increases after DSS and remains high 2 days of recovery. Lineage-traced Col4 expressing cells do not express the enteroendocrine (EE) marker Prospero, occasionally express the enterocyte marker Pdm1, and often die after repair is complete, related to Figure 2.**

**(A)** *Su(H)GBE>GFP* enteroblast cells were quantified in the posterior midgut, in untreated conditions, in vehicle-only conditions, in DSS conditions, or after recovery from DSS.

**(B)** Quantifications of the average number of *Su(H)GBE>mCD8GFP* colocalized with anti-Cleaved Caspase 3 post DSS. Shaded region represents the S.E.M.

**(C,D)** The complete data set from *Cg>GTRACE* experiments. *Cg>RFP* (magenta) with and without DSS, and *Cg*-lineage (green) with and without DSS, at various recovery time points. Each dot represents a posterior midgut.

**(E-G)** *Cg>RFP* cells (magenta) and *Cg*-lineage expressing cells (green). *Cg>G-TRACE* cells were not labeled with the EE cell marker anti-Prospero (E). 15% GFP+ *Cg>G-TRACE* cells labeled with the enterocyte marker anti-Pdm1 after 2 d recovery from DSS (50/340, example shown by arrow in G''') (F - No recovery, G - 2 d recovery). Scale bar, 10 µm.

**(H)** Quantifications of *Cg>RFP* or Cg-lineage cells labeled with anti-Cleaved Caspase 3 (CC3) with or without DSS.
