## Supplemental Figure 7 for "Piezo-dependent surveillance of matrix stiffness generates transient cells that repair the basement membrane"

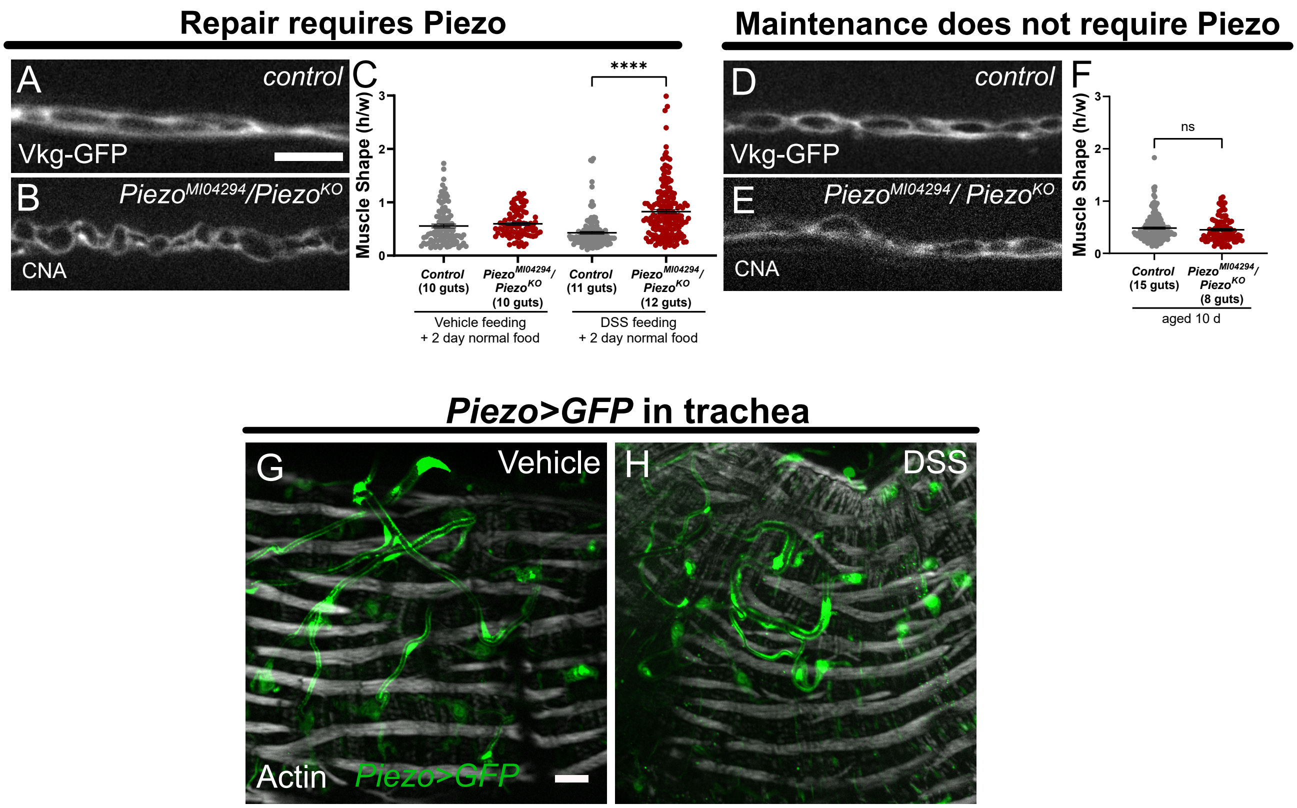


**Fig. S7. Specificity of Piezo mutation. Piezo is expressed in trachea, related to Figure 3.**

**(A-F)** Homozygous Piezo mutants (*Piezo*$^{MI04294}$/*Piezo*$^{KO}$) further demonstrate that *Piezo* is required for repair but not maintenance, assessed by the increased muscle shape following DSS and recovery. Basement membrane is visualized by Vkg-GFP in controls and the triple helix binding protein, CNA35, in the Piezo mutants. Mean +/- SEM, significance evaluated by *t*-test. Scale bar, 10 µm.

**(G,H)** Trachea of the posterior midgut express *Piezo-Gal4* (green) both with and without DSS. Tracheae are evident by their tubular morphology. Actin staining (gray) labels peristalsis muscles. Scale bar, 20 µm.
